# Spatial dynamics of feedback and feedforward regulation in cell lineages

**DOI:** 10.1101/2021.06.15.448623

**Authors:** Peter Uhl, John Lowengrub, Natalia Komarova, Dominik Wodarz

## Abstract

Feedback mechanisms within cell lineages are thought to be important for maintaining tissue homeostasis. Mathematical models that assume well-mixed cell populations, together with experimental data, have suggested that negative feedback from differentiated cells on the stem cell self-renewal probability can maintain a stable equilibrium and hence homeostasis. Cell lineage dynamics, however, are characterized by spatial structure, which can lead to different properties. Here, we investigate these dynamics using spatially explicit computational models, including cell division, differentiation, death, and migration / diffusion processes. According to these models, the negative feedback loop on stem cell self-renewal fails to maintain homeostasis, both under the assumption of strong spatial restrictions and fast migration / diffusion. Although homeostasis cannot be maintained, this feedback can regulate cell density and promote the formation of spatial structures in the model. Tissue homeostasis, however, can be achieved if spatially restricted negative feedback on self-renewal is combined with an experimentally documented spatial feedforward loop in which stem cells regulate the fate of transit amplifying cells. This indicates that the dynamics of feedback regulation in tissue cell lineages are more complex than previously thought, and that combinations of spatially explicit control mechanisms are likely instrumental.

## Introduction

Tissue homeostasis is central to the functioning and survival of higher organisms. Adult tissues are maintained by tissue stem cells that can both self-renew and differentiate to give rise to transit amplifying cells, which in turn give rise to terminally differentiated cells. Stem cells can divide asymmetrically [1], where one daughter is another stem cell and the other is a differentiating cell. In mammalian systems, however, data indicate that a stochastic symmetric division model also plays an important part, where a stem cell gives rise to either two daughter stem cells or to two daughter differentiated cells [2-4]. In such settings, it is thought that feedback mechanisms are required to prevent unbounded cell growth and to maintain homeostasis. Corresponding feedback signaling molecules have been much discussed in the literature in different tissues, and loss of feedback signals have been implicated in carcinogenesis [5, 6]. Several feedback molecules have been shown to determine the fate of cell divisions, influencing whether self-renewal or differentiation can occur. Examples include GDF11 and Activin βB, which negatively regulate self-renewal rates in progenitor and stem cells in the olfactory epithelium of mice [7, 8]; transforming growth factor beta (TGF-β) [9], which is mutated in a variety of tumors [10-12]; the bone morphogenetic protein 4 pathway (BMP4) that is inactivated in glioblastomas [13]; and the APC tumor suppressor gene that is inactivated in colorectal cancer, with concomitant activation of the Wnt cascade [14].

A growing mathematical literature has emerged that investigates feedback control in relation to tissue homeostasis, and loss of feedback control in relation to carcinogenesis [6-8, 15-26]. One particular approach focused on the notion that feedback factors produced by differentiated cells might play an important role for determining the fate of cell divisions. Specifically, in the olfactory epithelium, there appears to be negative feedback from differentiated cells both on the probability of stem cell self-renewal and on the rate of cell division, mediated by GDF11 and Activin βB [7, 8]. These observations motivated mathematical models showing that such negative feedback from differentiated cells onto stem cell division patterns can play an important role for the maintenance of tissue homeostasis [8, 24, 27] [21-23]. Mathematical models predicted that in the absence of this feedback, unbounded growth occurs, while negative feedback inhibiting stem cell self-renewal can result in a stable equilibrium.

These mathematical models of negative feedback regulation were all based on ordinary differential equations (ODEs), which assume perfect mixing of cells and molecules. In other words, no spatial structure was assumed. This applies to most mathematical models of stem cell regulation, with a few exceptions, e.g. [28]. Here, we re-formulate negative feedback models within cell lineages assuming spatially restricted dynamics, and specifically examine how this affects the ability of negative feedback loops to maintain tissue homoeostasis. We further construct models of other control mechanisms where spatial interactions are important and determine their effect on tissue homeostasis. We start by briefly reviewing the previously published ODE models that were used to study negative feedback from differentiated cells on stem cell division patterns in relation to the ability to maintain tissue homeostasis. We then introduce the spatial modeling framework and explore how this affects the dynamics.

### Basic models without spatial restriction

An ordinary differential equation model has been used to describe tissue hierarchy dynamics in a healthy tissue [8, 15, 24], and the models presented here build on these approaches. While cell lineages consist of stem cells, transit amplifying cells, and terminally differentiated cells, we can make a simplification and take into account only stem cells (which encompass all the proliferating cells) and differentiated cells [6]. Denoting stem cells (SC) by S and differentiated cells (DC) by D, the model is given by:

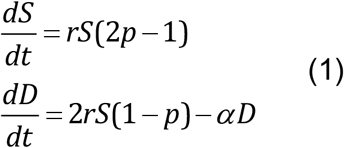

Stem cells divide with a rate r. With a probability p, the division results in two daughter stem cells (self-renewal), and with a probability (1-p), the division results in two daughter differentiated cells (differentiating division). Differentiated cells are assumed to die with a rate α. These equations capture a probabilistic model of tissue control, where on the cell population level a fraction of the symmetric divisions result in two daughter stem cells and the remaining fraction results in two daughter differentiated cells. In addition to symmetric divisions, asymmetric divisions may play a role in tissue renewal. With asymmetric cell division, a stem cell gives rise to one stem cell and one differentiated cell, thus maintaining a constant population of stem cells. While not included here, it has been previously shown that the incorporation of asymmetric cell divisions into this modeling framework does not fundamentally alter the properties of the model [29].

This system is only characterized by a neutrally stable family of nontrivial equilibria if p=0.5 [6, 8]. If p>0.5, infinite growth is observed. If p<0.5, the cell population goes extinct.

If we include the assumption that differentiated cells secrete negative feedback factors that influence stem cell division patterns, however, more realistic dynamics can be observed [6, 8, 24]. In particular, it has been shown that negative feedback from differentiated cells onto the probability of self-renewal, p, results in the existence of a stable equilibrium of cells, which might contribute to tissue homeostasis. This is because increased numbers of differentiated cells shift the division pattern in favor of differentiation, which limits overall cell growth. Mathematically, this has been expressed by p = p’/(1+f_1_D^κ1^), where p’ is the basic self-renewal probability of stem cells in the absence of any feedback. The existence of a stable equilibrium requires p’>0.5.

In addition, feedback loops have been proposed where differentiated cells reduce the rate of stem cell division [8, 24], which can be expressed as r = r’/(1+f_2_D^κ2^), where r’ is the basic stem cell division rate in the absence of any feedback. While this feedback can influence the dynamics of the system, it does not contribute to the existence of a stable equilibrium, and hence homeostasis.

Under these assumptions, the stable equilibrium population size is given by

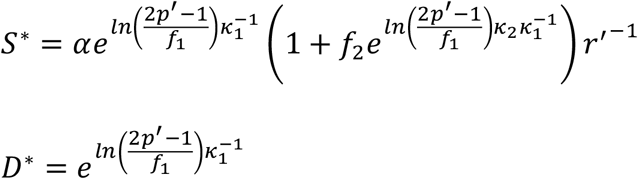

### An agent-based model with diffusion of feedback mediators

We consider a two-dimensional spatial model of growing cell populations that are regulated by a population of diffusing negative feedback mediators. The model contains a cell layer and a layer that describes the dynamics of feedback factors.

The dynamics of the cell population are described by a stochastic agent-based model, which assumes two cell populations: stem cells and differentiated cells. The agent-based model is given by a 2-dimensonal grid that contains nxn spots. Each spot can be empty, contain a stem cell, or contain a differentiated cell.

During one time step, the grid is sampled N times, where N is the total number of cells in the system. If the sampled spot contains a stem cell, a division event can occur with a probability p_div_. A target spot is chosen randomly from the eight nearest neighboring spots, into which the offspring cell can be placed. If that spot is already occupied, the division event is aborted. If the target spot is empty, a new stem cell is placed there with a probability p_self_ (self-renewal), and a differentiated cell is placed there with a probability 1-p_self_. A stem cell can die with a probability p_Sdeath_. If the sampled spot contains a differentiated cell, cell death occurs with a probability p_Ddeath_. In addition to these processes, stem and differentiated cells can attempt a migration event with a probability p_mig_. A spot is selected randomly from the eight nearest neighbors and if this spot is empty, the cell moves there.

The probability of self-renewal is influenced by the presence of feedback factors that can be secreted from differentiated cells. The dynamics of the feedback factors are described by a deterministic patch model. Each spot on the cell grid has a corresponding patch, in which the concentration of feedback factors is recorded. In each patch *i*, the concentration of the feedback factor, *z*_*i*_, is given by the following ODE: *dz*_*i*_*/dt = c – bz*_*i*_ *– mgz*_*i*_ *+ gZ*. The parameter *c* represents the production rate of the feedback factor. We set *c=0* if the spot does not contain a differentiated cell, otherwise *c>0*. Feedback factors are assumed to decay with a rate *b* in each patch. Feedback factors move to the nearest neighboring patches with a rate *g*, representing diffusion processes. The parameter *m* denotes the number of neighboring patches. Typically *m=8*, except for boundary patches, where *m<8*. The variable *Z* denotes the sum of all feedback factor populations, *z*_*i*_, among the directly neighboring patches. For each time step of the agent-based model, the ODEs in each patch were run for one time unit. The probability of self-renewal for a given cell in the agent-based part of the model is thus given by *p*_*self*_ *= p*^*(0)*^_*self*_ */ (1+hz*_*i*_), where *p*^*(0)*^_*self*_ denotes the self-renewal probability in the absence of any feedback. The more feedback factors are locally present in the patch corresponding to the spot in the cellular grid, the lower the probability of stem cell self-renewal (and the larger the probability of differentiation). The parameter *h* describes the strength of feedback inhibition, with larger values of *h* corresponding to more potent inhibition. For simplicity, we do not include feedback on the stem cell division rate, because this has been shown in the ODEs to not contribute to the existence of a stable equilibrium. This feedback can, however, be easily incorporated into the model in the same way.

The degree of spatial restriction in this system is given by two parameters, i.e. the cell migration rate of cells, p_mig_, and the rate of feedback diffusion, g. If p_mig_=0 and the value of g is low, the system is characterized by strong spatial restriction. For large values of p_mig_ and g, the system approaches mass action dynamics (perfect mixing).

As initial conditions, a square of 7×7 spots in the center of the grid was filled with stem cells, and the simulation was run according to the rules described above. Parameter values are largely unknown, and hence were chosen for the purpose of demonstration. Parameters quoted in the figures are scaled such that stem cells divide on average once a day [30, 31], and that terminally differentiated cells on average live for 10 days. The reported results, however, do not depend on these particular values.

### Dynamics assuming strong mixing of cells and feedback mediators

We start by assuming large values of *p*_*mig*_ and *g*, i.e. the migration rate of cells and the diffusion rate of feedback factors. In this limit, the average dynamics of the agent-based model converge to those predicted by a corresponding set of ordinary differential equations. Denoting the populations of stem cells, differentiated cells, and feedback factors by S, D, and Z, respectively, the ODEs are given as follows.

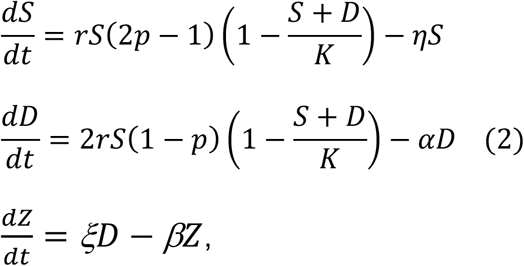

where negative feedback from differentiated cells onto the probability of self-renewal is given by 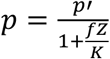, and p’ is the self-renewal probability of stem cells in the absence of feedback. In contrast to model (1) above, here we explicitly track the population of secreted feedback factors, Z. They are produced by differentiated cells with a rate ξ and decay with a rate β. Another change in the model (compared to model (1)) is the inclusion of a carrying capacity K, which corresponds to the maximum population size the system can sustain, independent of the negative feedback loop. This describes the finite grid size underlying the agent-based model. In the term describing the negative feedback on the stem cell self-renewal probability, the abundance of the secreted feedback mediator is divided by the carrying capacity K. The reason is that in the agent-based model, a larger system results in a lower amount of soluble feedback mediators available per cell due to diffusion, and this has to be captured in the corresponding ODE. Finally, we assumed that stem cells can die with a rate η, in addition to being removed through differentiation.

Figure 1A shows that this system of ODEs describes the average behavior of the agent-based model well for the limit of large cell migration and feedback diffusion rates. More generally, this ODE model is characterized by two equilibria. If p’<0.5, the cell population goes extinct and the system converges to the trivial equilibrium S^(0)^=0, D^(0)^=0, Z^(0)^=0. If p’>0.5, the system converges to a stable equilibrium at which all cell populations exist (not written down here due to complexity of expressions). An important property of this equilibrium is that the total cell population size is proportional to the carrying capacity, K (Figure 1B). As the value of the carrying capacity increases towards an infinitely large size, the number of cells also increase to an infinitely large size, despite the occurrence of negative feedback regulation. Therefore, in this model, negative feedback alone cannot maintain homeostasis of the cell population, defined by keeping the number of cells constant. The strength of negative feedback does, however, regulate the density of cells for a given carrying capacity, K. This behavior is in contrast to model (1) that does not take into account a carrying capacity. The reason is that model (1), as well as similarly structured models [24], assume that a given amount of feedback mediators is equally effective regardless of the system’s size, while model (2) assumes that the same amount of feedback mediators becomes less effective at suppressing stem cell self-renewal for larger system sizes, which corresponds to the formulation of the agent-based model and might be biologically more realistic.

**Figure 1.**
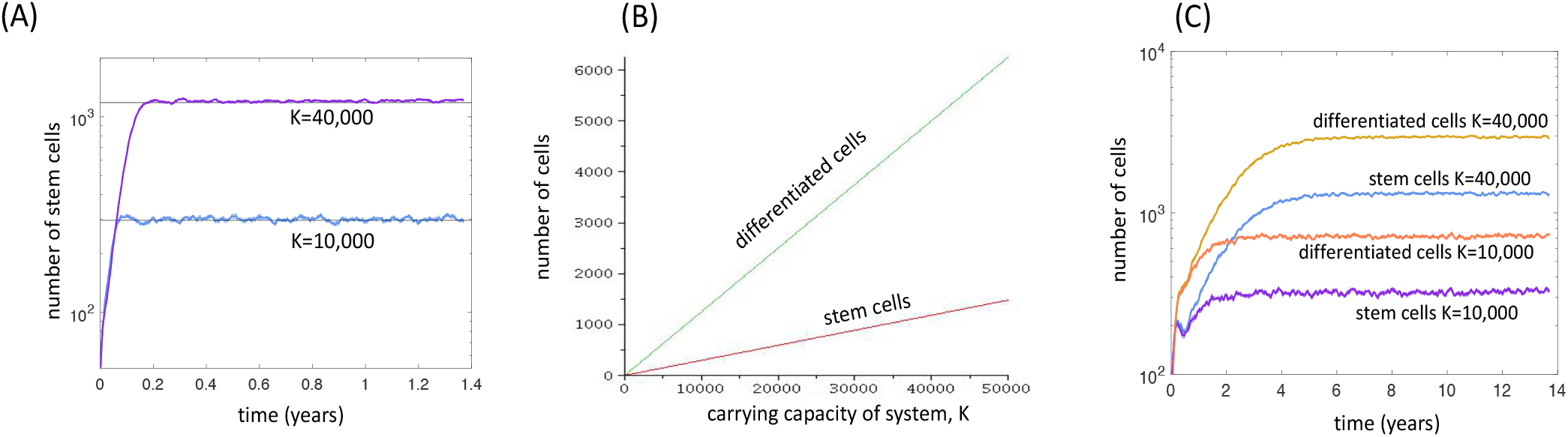
Properties of the spatially explicit computational model with negative feedback. (A) Dynamics of the agent-based model, depending the carrying capacity K under the assumption that cells migrate with a high rate (large p_mig_) and feedback mediators move across space at a high rate (large *g*). The straight lines represent equilibrium values derived from the corresponding ODE system (2). Parameters are given as follows. For agent-based model: P_div_=0.04167, p^(0)^_self_=0.7, P_Sdeath_=0, P_Ddeath_=0.0083, p_mig_=0.67, h=1.6, c=8.33, b=4.167, g=83.33; n=100 and n=200 for the small and large system, respectively. The average over 46 simulations are shown for each case; standard errors are too small to see. For ODEs: r=0.04167, η =0, α =0.0083, ξ =8.33, β =4.167, p’=0.7, f=1.6, K=100×100 and 200×200 for the small and large systems, respectively. (B) Equilibrium properties of the corresponding ODE system (2) as a function of the carrying capacity, K. Parameters were chosen as follows. r=0.04167, η=0, α =0.0083, ξ =8.33, β =4.167, p’=0.7, f=1.6. (C) Dynamics of the agent-based simulation, depending the carrying capacity K under the assumption of spatial restriction (p_mig_=0, low *g*). Parameters were chosen as follows. P_div_=0.04167, p^(0)^_self_=0.7, P_Sdeath_=0.000083, P_Ddeath_=0.004167, p_mig_=0., h=0.004, c=8.33, b=0.0083, g=0.833; n=100 and n=200 for the small and large system, respectively. The average over 46 simulations are shown for each case; standard errors are too small to see. Units of parameters are in hours.

### Dynamics assuming spatial restrictions

Here we assume that the processes of cell migration and the diffusion of feedback mediators are spatially restricted. For simplicity, we assume the strongest form of spatial restriction of cell migration, i.e. no migration occurs, and cells are assumed to leave their offspring in a nearest neighboring spot. For feedback mediators, the rate of diffusion will be varied from low to high. In general, similar properties are observed compared to the well-mixed system studied in the previous section. That is, negative feedback can only regulate the density of the cells, but not total numbers: The larger the size of the grid, the larger the total cell population size (Figure 1C). Beyond the properties of the well-mixed model, however, we find that negative feedback inhibition can result in pronounced spatial patterns, in which a number of islands of stem and differentiated cells exist, separated by space that is not occupied by cells (Figure 2A). As the strength of feedback inhibition is reduced (lower value of parameter h), the distribution of cells within the space becomes uniform (Figure 2B). We quantified the degree of clustering by calculating the ratio of the variance to mean of the number of cells (which we denote by σ) assuming the grid is subdivided into a number of relatively small squares. A ratio that is significantly greater than one indicates that the cell population is clustered across space. Figure 2C shows that as the strength of negative feedback inhibition is reduced, there is a sharp transition from a clustered distribution of cells towards a uniform distribution. If the clustered population structure corresponds to a normal state, then it is possible that the transition to a uniform distribution due to reduced negative feedback corresponds to a first step towards abnormal growth. Interestingly, in the presence of strong negative feedback and clustered population structure, the stem cell population is in the minority (Figure 2). For weak negative feedback and uniform cell distribution, however, the stem cell fractions are significantly larger (Figure 2).

**Figure 2.**
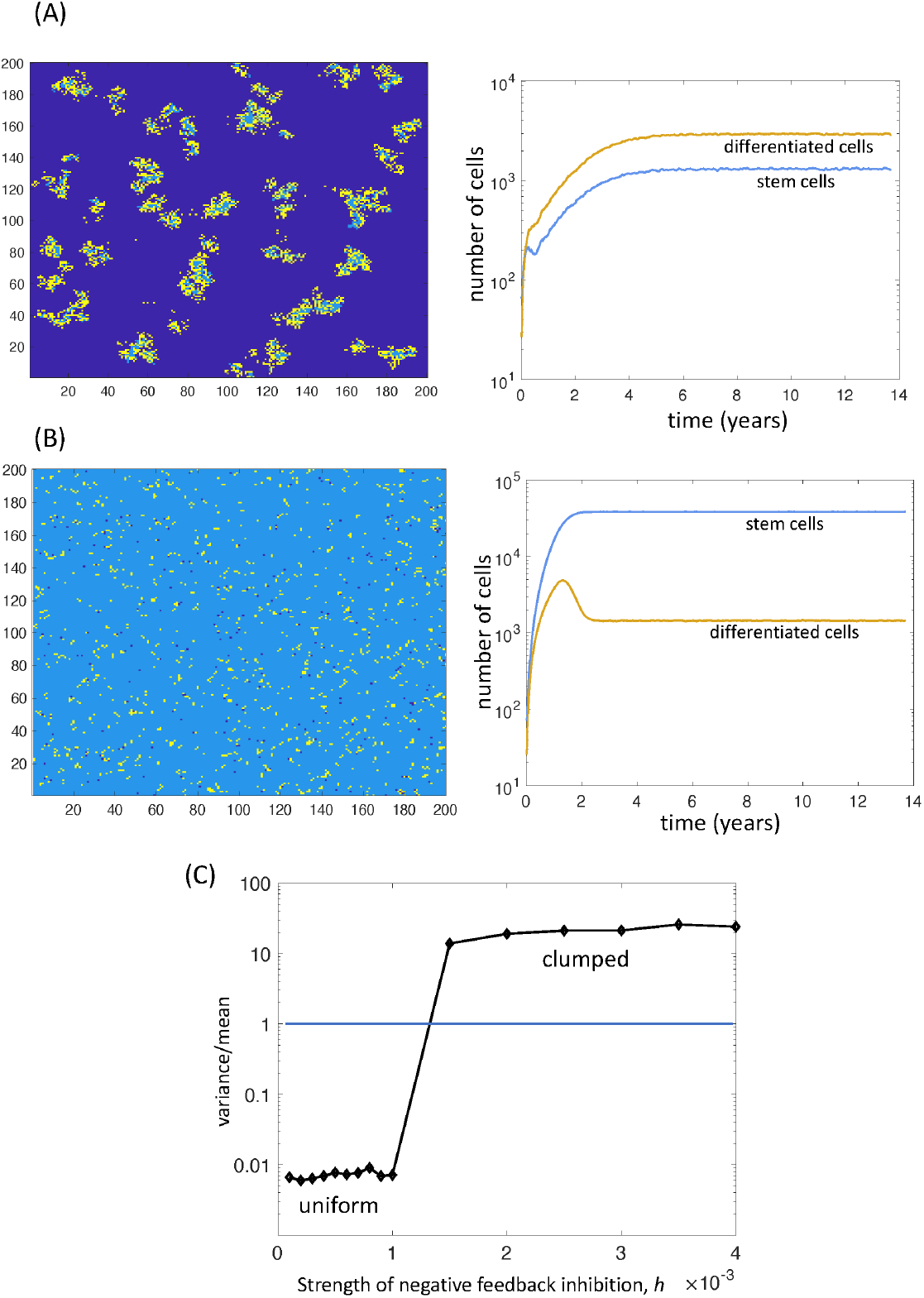
Spatial patterns observed in the agent-based model with negative feedback. Dark blue is empty space, light blue represents stem cells, and yellow represents differentiated cells. (A) With a stronger degree of negative feedback on the stem cell self-renewal rate, clumped spatial patterns are observed. Islands of stem and differentiated cells form, separated by empty space. Stem cells are in the minority. The spatial picture is a snapshot in time at equilibrium, and the time series represents the average over 46 simulations; standard errors are too small to see. (B) For weaker negative feedback, these spatial patterns break down, and a uniform distribution of cells across space is observed. Also, stem cells become the dominant population. The spatial picture is a snapshot in time at equilibrium, and the time series represents the average over 46 simulations; standard errors are too small to see. (C) The degree of clumpiness on the distribution of cells across space can be quantified by dividing the space up into relatively small squares, and the number of cells per square is recorded. If the ratio of σ= variance / mean is greater than 1, the spatial pattern is clumped. If the ratio σ is less than one, the distribution is uniform. The graph shows the value of σ at the end of the simulation (at equilibrium). As the rate of negative feedback inhibition is increased from low to high, we observe a relatively sharp transition in the ratio σ, i.e. from a uniform to a clumped distribution of cells across the space. Baseline parameter values were chosen as follows. P_div_=0.04167, p^(0)^_self_=0.7, P_Sdeath_=0.000083, P_Ddeath_=0.004167, p_mig_=0., h=0.004, c=8.33, b=0.0083, g=0.833; n=200. For (A) h=0.004, (B) h=0.001, and for (C) the value of h was varied, as indicated. Units of parameters are in hours.

Finally, we note that if the strength of negative feedback inhibition is large and crosses a threshold in this model, extinction of the cell population is observed.

Next we investigated how the different outcomes depend on model parameters. This was done by randomly drawing the logarithm of parameter values from a uniform distribution and recording the ratio σ. In Figure 3, two parameters were varied simultaneously for any given plot: The rate of negative feedback inhibition on stem cell self-renewal, *h*, was always varied, together with a second parameter. The outcome of the simulation is color-coded in Figure 3, in which each dot represents the outcome of an individual simulation in the parameter space. A value of σ>1.5 (clustering) is recorded in red, and a value of σ<1.5 (uniform distribution) is recorded in blue. Runs in which population extinction occurred are shown in yellow. As mentioned above, clustered cell population persistence occurs for intermediate rates of negative feedback inhibition, while larger and smaller rates of feedback inhibition result in population extinction, or a uniform distribution of cells across space, respectively. The width of the parameter region in which cell clusters are observed depends on parameters (Figure 3). In particular, it depends on the diffusion rate of feedback mediators, and on the decay rate of the feedback mediators. Larger diffusion rates, *g*, and slower decay rates of feedback mediators, *b*, lead to a broader region of feedback inhibition rates for which cell clusters are observed (Figure 3). If the diffusion rate of feedback mediators is slow and their decay rate is fast, the mediators secreted by a given cell mostly act locally, and the clustering regime is narrow. In the limit, the clustering regime is so narrow that the behavior of the system essentially transitions from extinction to a uniform invasion of the space by cells (Figure 3). If the diffusion rate of feedback mediators is faster and they decay slower, then the mediators secreted from a given cell affect cells in a larger area of the space, and the clustering behavior becomes pronounced (Figure 3). Therefore, the model suggests that the clustered persistence of cells requires feedback mediators to act beyond the immediate neighborhood of the cell from which they are secreted. Other model parameters do not have a significant influence on the range of feedback inhibition values across which clustered cell persistence is observed (Figure 3).

**Figure 3.**
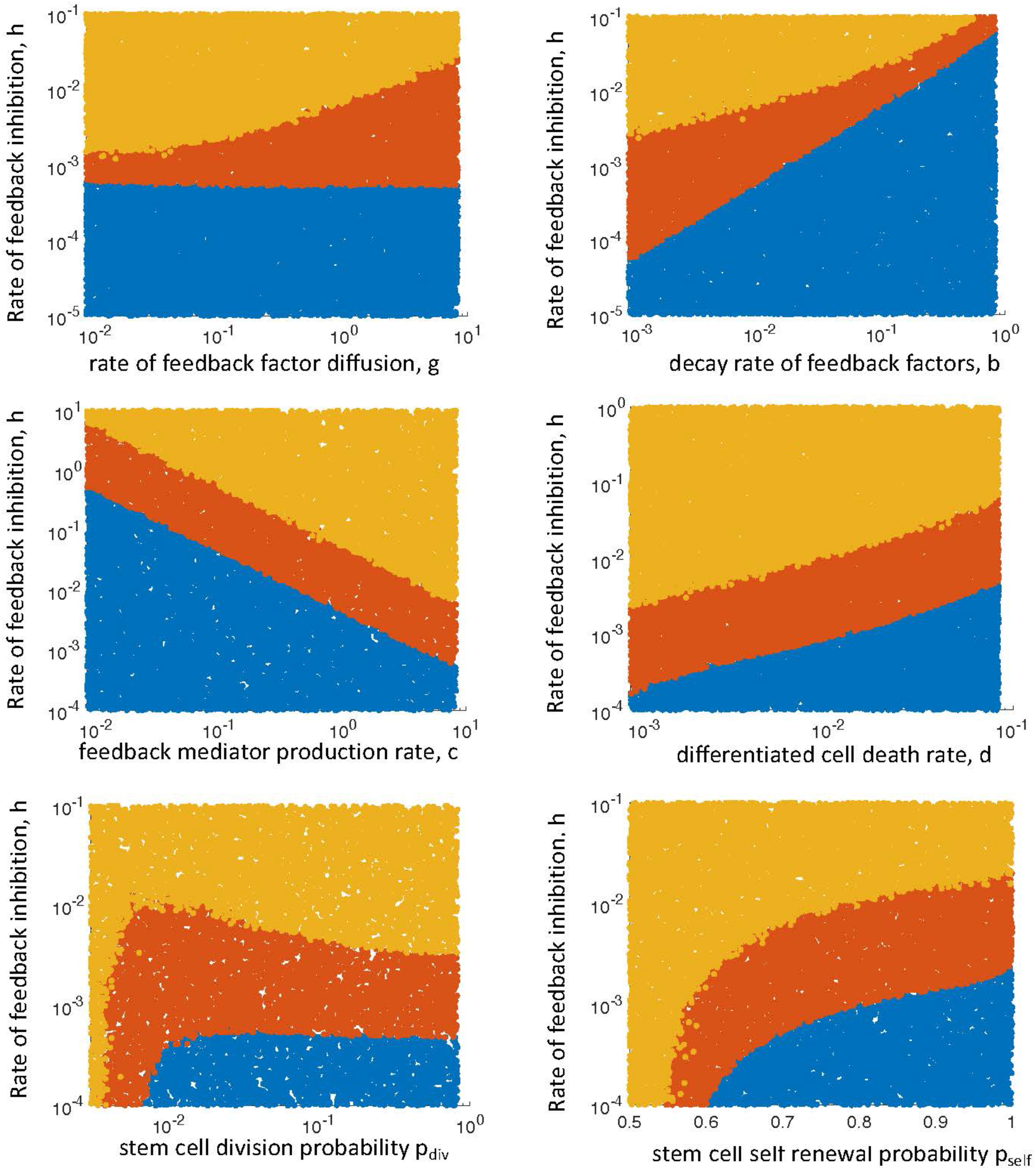
Parameter dependencies of outcome. Each dot in the graph represents the long-term outcome of an individual simulation. Each simulation was run up to a time threshold, and the spatial distribution was determined by calculating σ = variance / mean. Yellow indicates the extinction of the cells. Blue indicates a distribution that is characterized by σ < 1.5 (mostly uniform distribution). Red indicates σ > 1.5 (clumped distribution of cells). For each graph, two parameters were varied: the strength of negative feedback on the stem cell self-renewal probability, h, and a second parameter, as indicated in the individual graphs. Baseline parameters were chosen as follows. P_div_=0.04167, p^(0)^_self_=0.7, P_Sdeath_=0.000083, P_Ddeath_=0.004167, p_mig_=0., c=8.33, b=0.0083, g=0.833; n=200. Units of parameters are in hours.

### Feedforward loop

So far, we have considered regulatory loops where differentiated cells influence the behavior of stem cells. Data from the airway epithelial tissue from mice [32], however indicate that stem cells can also send a signal forward to their progeny and influence their behavior. In fact, this “feedforward” regulation is inherently spatial. The stem cells secrete a notch ligand to their daughter transit amplifying cells (secretory cells), and this signal is necessary to maintain the transit amplifying cell population. In the absence of this signal, the transit amplifying cells undergo terminal differentiation to become ciliated cells. Hence, the further the transit amplifying cells are located away from stem cells, the weaker this signal, and the more likely terminal differentiation occurs. Transit amplifying cells that are located closer to stem cells receive a stronger signal and are more likely to be maintained.

We adapt our above described agent-based model to describe this scenario. To do so, we expand the model complexity to include a population of transit amplifying cells (TA) in addition to stem cells. Now, stem cell differentiation results in the generation of two TA cells. Similarly to stem cells, TA cells have a probability to divide (q_div_). With a probability q_self_, this is a self-renewing division, giving rise to two TA daughter cells; with a probability 1-q_self_, this is a terminally differentiating division, giving rise to two daughter differentiated cells. TA cells are assumed to die with a probability q_Tdeath_. Feedforward mediators are secreted from stem cells. The dynamics of the feedforward factors are again described by a deterministic patch model. Each spot on the cell grid has a corresponding patch, in which the concentration of feedforward factors is recorded. In each patch *i*, the concentration of the feedforward factor, *w*_*i*_, is given by the following ODE: *dw*_*i*_*/dt = c*_*2*_ *– b*_*2*_*w*_*i*_ *– mg*_*2*_*w*_*i*_ *+ g*_*2*_*W*. The parameter *c*_*2*_ represents the production rate of the feedback factor. We set *c*_*2*_*=0* if the spot does not contain a stem cell, otherwise *c*_*2*_*>0*. Feedforward factors are assumed to decay with a rate *b*_*2*_ in each patch. Feedforward factors move to the nearest neighboring patches with a rate *g_2_*, representing diffusion processes. The parameter *m* again denotes the number of neighboring patches. The variable *W* denotes the sum of all feedforward factor populations, *w*_*i*_, among the directly neighboring patches. Importantly, the probability of TA cell self-renewal (and thus maintenance) is determined by the concentration of the feedforward factor according to 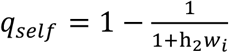. For large feedforward factor concentrations, the probability of TA self-renewal converges to one.

In the absence of feedforward factors, the probability of TA self-renewal is zero. In other words, the feedforward factors secreted by stem cells are responsible for the maintenance of the TA cell population, and in the absence of these mediators, terminal differentiation is the only fate of the division.

It is instructive to start examining the dynamics of this system under the assumptions that stem cells and TA cells do not die (and only disappear through differentiation). In this case, a structure forms where stem cells are located in the center, surrounded by an area of TA cells, with differentiated cells at the periphery (Figure 4Ai). Interestingly, this structure is self-contained and stops growing after a while. The size of this structure is independent of the grid size, and determined by the diffusion rate of the feedforward factor. The further the feedforward factors can diffuse from the stem cells, the larger the area of the TA cell population. In other words, in this scenario, the feedforward regulation can maintain homeostasis of cell numbers, and the same cell numbers are maintained no matter how large the available space is. Hence, this is true homeostasis. The reason for this behavior is that there is competition between stem and TA cells for space, similar to the dynamics described by Hillen et al [33]. Stem cells enable self-renewal of TA cells in their immediate vicinity, and the TA cells block the stem cells from dividing further. Hence, the growth of the overall cell population is limited.

**Figure 4.**
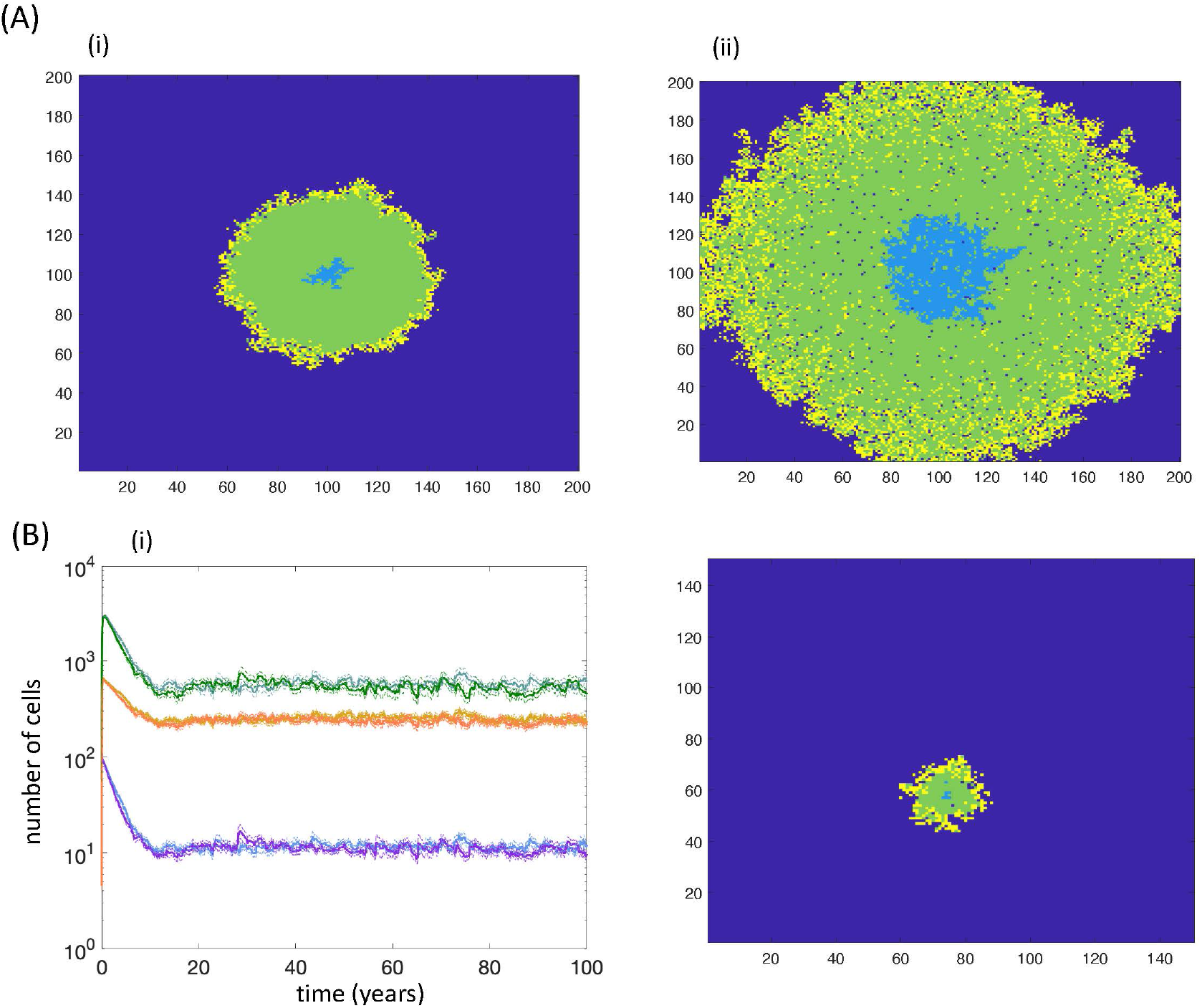
(A) Properties of the agent-based models with the feedforward control loop only. Dark blue represents empty space, light blue stem cells, green transit amplifying cells, and yellow terminally differentiated cells. (i) In the absence of stem and transit amplifying cell death, the cell population stops growing and converges to an equilibrium that is independent of the carrying capacity, K (not shown). The transit amplifying cells block stem cell divisions due to lack of available space, and this prevents the area of cells from expanding. The picture shown corresponds to the population at steady state. Parameters were as follows: P_div_=0.04167, p^(0)^_self_=0.8, q_div_=0.0583, P_Sdeath_=0, P_Tdeath_=0, P_Ddeath_=0.004167, p_mig_=0., c_2_=0.833, b_2_=0.0083, g_2_=0.4167, h_2_=2, n=200. (ii) In the presence of cell death, however, this mechanism breaks down and the stem cells can continuously expand into empty space, provided by the death of transit amplifying cells. The picture represents a snap-shot during this cell expansion. Parameters were chosen as follows: P_div_=0.04167, p^(0)^_self_=0.8, q_div_=0.0583, P_Sdeath_=0.000083, P_Tdeath_=0.0001, P_Ddeath_=0.004167, p_mig_=0., c_2_=0.833, b_2_=0.0083, g_2_=0.4167, h_2_=2, n=200. (B) Properties of the agent-based model that contains both the feedforward and the feedback loop, and assumes the occurrence of death for all cell populations. (i) Time series, in which the system / grid size was varied, n=100 vs n=150. Blue and purple show stem cells for the smaller and larger grid size, respectively. Light green and dark green show TA cells, for the smaller and larger grid size, respectively. Yellow and orange show differentiated cells, for the smaller and larger grid size, respectively. The lines present the average time series over 46 iterations of the simulation, and the dashed lines represent the average plus minus standard errors. (ii) A snapshot of the spatial configuration of cells at a specific time point during steady state. Parameters were chosen as follows. P_div_=0.04167, p^(0)^_self_=0.8, q_div_=0.0583, P_Sdeath_=0.000083, P_Tdeath_=0.0001, P_Ddeath_=0.004167, p_mig_=0., c=8.33, b=0.0833, g=0, h=0.06, c_2_=8.33, b_2_=0.0833, g_2_=3.33, h_2_=2.5. Units of parameters are in hours.

This homeostasis, however, represents a somewhat artificial situation due to the assumption that stem and TA cells can only be eliminated through differentiation processes, and no explicit cell death is assumed to occur. If cell death is assumed to occur in stem and TA cells, the spatial competition dynamics either go in favor of the TA cells, resulting in the exclusion of stem cells and thus in the extinction of the whole cell population, or in favor of stem cells, in which case the stem cell population can expand outward over time, leading to cell population growth limited by the available space (Figure 4Aii), similar to the simulations with negative feedback on stem cell self-renewal. Hence, the homeostasis observed in this system is lost in the presence of stem cell death.

### Combination of negative feedback and feedforward regulation

The above sections considered negative feedback on the stem cell self-renewal probability, and feedforward regulation separately. For each model, these mechanisms failed to maintain homeostasis of cell numbers. Here, we consider a modified version of the agent-based model that includes both of these control processes at the same time. The two control mechanisms are implemented in the same way as described above. We now observe parameter regions in which true homeostasis is maintained (Figure 4B), i.e. the number of cells converges to an equilibrium value that is independent of the size of the grid or carrying capacity (including parameter regions where stem and transit amplifying cells are assumed to die). The reason for this behavior is as follows. With the feedforward loop, the death of transit amplifying cells can make space for the invasion of self-renewing stem cells. If only the feedforward loop exists, this results in a continuous expansion of the stem cell population through space, which results in a concomitant expansion of transit amplifying and differentiated cells, and hence in uncontrolled growth. The addition of negative feedback on stem cell self-renewal, however, counters this process. The negative feedback mediators are assumed to be secreted from differentiated cells, which are located predominantly at the surface of the cell mass (Figure 4B). As the stem cells expand in space and reach locations closer to the differentiated cells, the negative feedback loop becomes important and counters this outward stem cell expansion by forcing terminal differentiation to occur. This prevents the unbounded expansion of the stem cells and ensures the existence of an equilibrium that is independent of the grid size (carrying capacity). This effect is observed even if the negative feedback factors act predominantly on a local level, including with zero diffusion rates (as long as the feedback mediators remain present for long enough to stop an expanding front of stem cells).

Exploration of the parameter space is computationally not feasible on a larger scale because each parameter combination needs to be run many times to obtain the average trajectories and thus to determine whether grid size determines the equilibrium number of cells. Additional simulations, however, are shown in the Supplementary Materials demonstrating that these dynamics are observed over varying parameter ranges.

It is important to point out that homeostasis is a result of spatial dynamics. It is possible to describe a non-spatial version of the feedforward control mechanism, where stem cells promote the self-renewal of TA cells regardless of spatial location. A corresponding ODE model that takes into account both the feedforward and the negative feedback mechanisms is presented in the Supplementary Materials. In the absence of space, the ability to of these controls to maintain homeostasis is lost, and the equilibrium cell population sizes are directly proportional to the carrying capacity of the system.

## Discussion

Previous mathematical modeling approaches [6, 8, 24, 27], based on the assumption that cells mix perfectly (mass action), suggested that negative feedback from differentiated cells on the self-renewal probability of stem cells is an important determinant of tissue homeostasis. Here, we extended this analysis into a spatially explicit scenario where cells grow in a finite space, characterized by a given carrying capacity (maximum possible number of cells the system can sustain). This suggests that the negative feedback from differentiated cells onto the stem cell self-renewal probability cannot by itself maintain tissue homeostasis. The number of cells at equilibrium always scales with the carrying capacity of the system, even under the assumption that cells and feedback mediators mix relatively well due to fast migration and diffusion processes. In these models, an infinitely large space available for cell growth will lead to infinitely large cell population sizes, despite the presence of the negative feedback loop.

The negative feedback on stem cell self-renewal can only maintain cell density, not total cell numbers in the models considered here. In the spatially explicit model versions (without significant migration of cells), this negative feedback can further lead to the formation of spatial patterns. In the presence of relatively strong negative feedback, clumps of cells (containing both stem and differentiated cells) form in the model, separated by empty space. This is interesting, because it suggests that this negative feedback loop might be involved in the formation of spatial structures in the cellular microenvironment. At the same time, however, the formation of spatial structures around stem cell populations, in particular the formation of stem cell niches, is highly complex and includes many components not currently taken into account in our model [34]. As the strength of feedback inhibition is reduced, there is a sharp transition in the model away from the clumped spatial structure towards a uniform distribution of cells, where the stem cell population is dominant. This means that loss of negative feedback on stem cell self-renewal might lead to a collapse of spatial organization, which might be a first step towards malignancy, even though uncontrolled growth does not yet occur. Whether this is indeed the case requires further investigation, in particular using models that take into account a higher biological complexity that characterize stem cell niche morphology.

While the model suggests that negative feedback on stem cell self-renewal can lead to spatial structures, the model further indicates that stronger spatial restrictions of feedback mediator diffusion limits the parameter regime in which this behavior is observed. The pattern formation is observed over relatively wide parameter regions only if the rate of feedback mediator diffusion is fast, i.e. if feedback factors secreted by a given cell can affect cells in a location further away. If feedback mediators are assumed to act only locally (through limited diffusion), the parameter region in which the spatial patterns form becomes vanishingly small. In this case, as the strength of the negative feedback is reduced from high to low, the model behavior more or less transitions from population extinction (stronger feedback) to a uniform distribution of cells (weaker feedback).

It was interesting to observe, however, that a combination of the negative feedback loop with a feedforward loop from stem cells to transit amplifying cells can lead to true homeostasis, where cell numbers settle around an equilibrium that is independent of the amount of space available (carrying capacity). The feedforward loop assumed in the model was based on data from the airway epithelium in mice, where stem cells were shown to secrete a notch ligand, which enabled the maintenance of the transit amplifying cells [32]. In the absence of this feedforward signal, transit amplifying cells were shown to undergo terminal differentiation. This is an inherently spatial process, since the concentration of the feedforward mediator decreases with distance from the originating stem cell. This leads to the formation of a structure, where stem cells are located in the center, surrounded by transit amplifying cells, while differentiated cells are located at the surface of this area. The presence of only the feedforward control loop cannot maintain homeostasis because the stem cell population can expand outward and replace transit amplifying cells. The concomitant presence of the negative feedback loop, however, results in increased differentiation (rather than self-renewal) of stem cells, limiting their ability to spread outwards in space. This contributes to a stable equilibrium that is independent of the carrying capacity of the system. This type of equilibrium is observed even if the diffusion rate of the negative feedback mediator is assumed to be slow, i.e. if the negative feedback loop acts largely on a local level. Importantly, an equivalent non-spatial model was not characterized by control-mediated cell homeostasis. These results suggest that the interplay between different feedback control mechanisms within a cell lineage in a spatial setting can lead to true homeostasis, which we could not observe in a model without spatial structure.

While the combination of these particular feedback and feedforward mechanisms can maintain true homeostasis in our model, it is likely that other feedback configurations exist that give rise to similar results. In addition, it is important to point out that this is a relatively simple model that was aimed to test how previously described homeostatic mechanisms [6, 8, 24] translate into spatial settings. As mentioned above, additional biological complexities, especially those that characterize stem cell niches, need to be considered to gain a more detailed picture of cell dynamics under homeostatic conditions. Besides the control loops involved in cells of a given lineage, complex signaling mechanisms between the cell lineage and its microenvironmental components exist that together yield homeostatic properties. These processes will need to be decoded with a combination of experiments and dynamical models. The results described here form a basis for understanding feedback dynamics in such more complex and realistic settings.

