## Supplementary Materials for "Spatial dynamics of feedback and feedforward regulation in cell lineages"

### ODEs corresponding to the agent-based model with feedback and feedforward control under fast mixing assumptions

Under fast migration of cells and fast diffusion rates of feedback and feedforward mediators, the dynamics in the agent-based model (main text) can be approximated by the following set of ODEs:

$$\frac{dS}{dt} = rS(2p-1)\left(1 - \frac{S+T+D}{K}\right) - \eta S$$

$$\frac{dT}{dt} = 2rs(1-p)\left(1 - \frac{S+T+D}{K}\right) + \rho T(2q-1)\left(1 - \frac{S+T+D}{K}\right) - \gamma T$$

$$\frac{dD}{dt} = 2\rho T(1-q)\left(1 - \frac{S+T+D}{K}\right) - \alpha D$$

$$\frac{dZ}{dt} = \xi_1 D - \beta_1 Z$$

$$\frac{dW}{dt} = \xi_2 S - \beta_2 W$$

The equations build on model (2) in the main text, and include stem cells, S, transit amplifying cells, T, and terminally differentiated cells, D. Differentiation of stem cells results in the generation of transit amplifying cells. Transit amplifying cells can divide with a rate $\rho$ and die with a rate $\gamma$. The division results in self-renewal with a probability q and in further differentiation with a probability 1-q, giving rise to terminally differentiated cells. The feedback factors are denoted by Z and their dynamics are described in the same way as in model (2) in the main text. Thus, the probability of self-renewal is again given by $p = \frac{p'}{1+\frac{f_1 Z}{K}}$, where p' is the self-renewal probability of stem cells in the absence of feedback. The feedforward factors are denoted by W. They are produced by stem cells with a rate $\xi_2$, and decay with a rate $\beta_2$. Their presence results in a higher probability of transit amplifying cell self-renewal, i.e. $q = 1 - \frac{1}{1+f_2 W/K}$.

It is possible to show that the equilibrium values (if any) of all the variables in the system of equations discussed here, are proportional to the carrying capacity, K. To see this, we divide all the equations by K and introduce a new, rescaled set of variables: $S_0=S/K$, $T_0=T/K$, $D_0=D/K$, $Z_0=Z/K$, and $W_0=W/K$. Then we can see that, when rewritten in terms of the rescaled variables, the system no longer contains parameter K (this includes the nonlinear terms containing probabilities p and q). As a consequence, the equilibrium values of quantities $S_0$, $T_0$, $D_0$, $Z_0$, and $W_0$ (if they exist) are K-independent, while the original variables have equilibrium values proportional to K. More generally, this argument shows that all the solutions of the system scale with the carrying capacity, K. Therefore, true homeostasis of cell numbers that is independent of system size, as was observed in the corresponding spatial version of this model, is not observed here.

### Spatial dynamics in the presence of feedback and feedforward control loops: Variations in parameters

The main text has shown that true homeostasis in cell numbers can be observed in the spatial agent-based model that includes both the feedback and the feedforward control loops. This was demonstrated with computer simulations under a particular combination of parameters. Figure S1 shows similar outcomes under different parameters. A comprehensive and systematic exploration of the parameter space was not possible due to prohibitive computational costs.

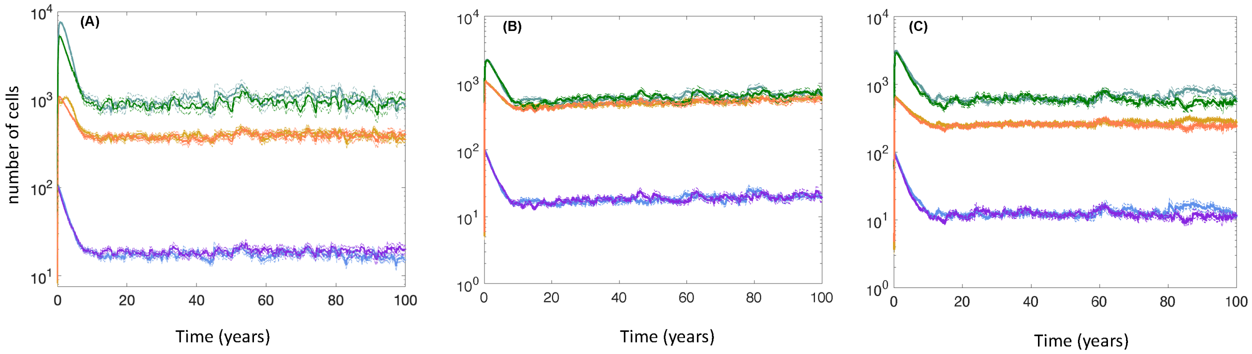


**Figure S1.** Same kind of simulation as Figure 4Bi, but with different parameter combinations, under which true cell homeostasis is observed that is independent of the system size. (A) $P_{div}=0.04167$, $p^{(0)}_{self}=0.8$, $q_{div}=0.0583$, $P_{Sdeath}=0.000083$, $P_{Tdeath}=0.0001$, $P_{Ddeath}=0.004167$, $p_{mig}=0.$, c=0.833, b=0.00833, g=0.04167, h=0.01, $c_2=0.833$, $b_2=0.00833$, $g_2=1.25$, $h_2=3$. (B) $P_{div}=0.04167$, $p^{(0)}_{self}=0.8$, $q_{div}=0.0583$, $P_{Sdeath}=0.000083$, $P_{Tdeath}=0.0001$, $P_{Ddeath}=0.00083$, $p_{mig}=0.$, c=8.33, b=0.083, g=0., h=0.01, $c_2=8.33$, $b_2=0.083$, $g_2=4$, $h_2=2$. (C) $P_{div}=0.04167$, $p^{(0)}_{self}=0.8$, $q_{div}=0.0583$, $P_{Sdeath}=0.000083$, $P_{Tdeath}=0.0001$, $P_{Ddeath}=0.004167$, $p_{mig}=0.$, c=8.33, b=0.083, g=0., h=0.02, $c_2=8.33$, $b_2=0.083$, $g_2=4$, $h_2=2.5$. Units of parameters are in hours. In all plots, the two grid sizes were n=100 and n=150, respectively. Blue and purple show stem cells for the smaller and larger grid size, respectively. Light green and dark green show TA cells, for the smaller and larger grid size, respectively. Yellow and orange show differentiated cells, for the smaller and larger grid size, respectively. The lines present the average time series over 46 iterations of the simulation, and the dashed lines represent the average plus minus standard errors.
